# Structural basis of interdomain communication in PPARγ

**DOI:** 10.1101/2022.07.13.499031

**Authors:** Sarah A. Mosure, Paola Munoz-Tello, Kuang-Ting Kuo, Brian MacTavish, Xiaoyu Yu, Daniel Scholl, Christopher C. Williams, Timothy S. Strutzenberg, Jess Li, Jared Bass, Richard Brust, Eric Kalkhoven, Ashok A. Deniz, Patrick R. Griffin, Douglas J. Kojetin

## Abstract

The nuclear receptor peroxisome proliferator-activated receptor gamma (PPARγ) regulates transcription via two activation function (AF) regulatory domains: a ligand-dependent AF-2 coregulator interaction surface within the C-terminal ligand-binding domain (LBD), and an N-terminal disordered AF-1 domain (NTD or A/B region) that functions through poorly understood structural mechanisms. Here, we show the PPARγ AF-1 contains an evolutionary conserved Trp-Pro motif that undergoes *cis/trans* isomerization, populating two long-lived conformations that participate in intradomain AF-1 and interdomain interactions including two surfaces in the C-terminal LBD (β-sheet and the AF-2 surface), which are predicted in AlphaFold 3 models but not AlphaFold 2. NMR and chemical crosslinking mass spectrometry show that interdomain interactions occur for soluble isolated AF-1 and LBD proteins, as well as in full-length PPARγ in a phase separated state. Mutation of the region containing the Trp-Pro motif, which abrogates *cis/trans* isomerization of this region, impacts LBD interaction and reduces basal PPARγ-mediated transcription and agonist-dependent activation of PPARγ. Our findings provide structural insight into published *in vitro* and cellular studies that reported interdomain functional communication between the PPARγ AF-1 and LBD suggesting some of these effects may be mediated via AF-1/LBD interactions.

## INTRODUCTION

Peroxisome proliferator-activated receptor gamma (PPARγ NR1C3) is a nuclear receptor transcription factor that controls gene expression programs influencing the differentiation of mesenchymal stem cells into adipocytes (adipogenesis), lipid metabolism, and insulin sensitivity. Like other nuclear receptors ^1^, PPARγ is a multidomain protein that contains a central DNA-binding domain (DBD) flanked by two regulatory regions that influence transcription: an N-terminal ligand-independent AF-1 domain (also called the NTD or A/B region) and a C-terminal ligand-dependent AF-2 coregulator interaction surface within the C-terminal LBD.

The molecular and structural basis of ligand-regulated functions of the LBD of nuclear receptors are relatively well understood. For PPARγ, structural biology studies including X-ray crystallography, hydrogen-deuterium mass spectrometry (HDX-MS), chemical crosslinking MS (XL-MS), and NMR spectroscopy have revealed how agonist ligands stabilize a transcriptionally active AF-2/helix 12 surface conformation upon binding to the orthosteric ligand-binding pocket in the LBD to promote coactivator protein recruitment and increased expression of PPARγ target genes that drive adipogenesis ^2–6^. More recently, a structural mechanism of ligand-dependent corepressor-selective PPARγ inverse agonism was reported ^7^. PPARγ inverse agonists enabled a structural definition of a transcriptionally repressive AF-2/helix 12 conformation that promotes corepressor interaction and transcriptional repression of PPARγ and a structural characterization of the apo-LBD conformational ensemble, which dynamically exchanges between active- and repressive-like conformations ^8,9^.

Despite these and other key advances in determining ligand-dependent structural mechanisms of nuclear receptor LBD function, the structural basis by which the disordered N-terminal AF-1 influences the function of PPARγ and other nuclear receptors remains poorly understood. Only a few crystal structures of full-length nuclear receptors including PPARγ have been reported and these structures either lack electron density for the disordered AF-1 or the AF-1 was removed to facilitate crystallization ^10–13^. Cryo-EM studies of nuclear receptors thus far have only provided low resolution (>10–25Å) structural snapshots of the AF-1 or, similar to the crystallography studies, protein samples were used where the AF-1 was removed ^14–17^. Furthermore, AlphaFold ^18^ models of nuclear receptors frequently show cloud-like AF-1/NTD structural depictions that are thought to be an artifact of the computational method to avoid steric clashes with structured domains ^19^.

Obtaining atomic resolution structural data on the PPARγ AF-1 would be important for the field—new regulatory mechanisms are likely to emerge, and the data may explain published observations of interdomain functional communication between the PPARγ AF-1 and AF-2/ LBD. For example, PPARγ-mediated transcription and adipogenesis is increased by AF-1 removal, indicating the AF-1 negatively regulates PPARγ-mediated transcription ^20^. Phosphorylation of Ser112 within the AF-1 negatively affects LBD functions (ligand binding and coregulator interaction), downregulates the expression of PPARγ target genes, and inhibits adipogenesis ^21–24^. Furthermore, phosphorylation of Ser112 in the AF-1 is inhibited by an agonist ligand binding to the LBD that stabilizes a transcriptionally active AF-2 surface conformation, but not by an inverse agonist that stabilizes a transcriptionally repressive LBD conformation ^25^. These observations suggest that the N-terminal AF-1 somehow alters the structure and function of the C-terminal LBD, and vice versa, though currently there is no structural evidence into the molecular mechanisms.

Here, we used biophysical and structural biology approaches suitable for studying intrinsically disordered proteins (IDPs) including ^1^H- and ^13^C-detected NMR spectroscopy, single-molecule FRET (smFRET), and XL-MS to gain molecular insight into how the disordered PPARγ AF-1 domain. Our studies uncovered several previously unknown structural features of the AF-1 that are poised to regulate the structure and function of PPARγ. Although the AF-1 is structurally disordered, our studies reveal intradomain contacts within the AF-1 indicating it adopts a partially compact conformational ensemble regulated by negative charge repulsion. We also uncovered a region of the AF-1 that natively exchanges between two long-lived structural conformations resulting from proline *cis/trans* isomerization at an evolutionarily conserved Trp-Pro motif. Finally, we show that the LBD physically interacts with the AF-1 and where the slowly exchanging AF-1 Trp-Pro motif interacts with two LBD surfaces including the AF-2. Our findings provide a structural rationale to explain previous reports of interdomain functional communication and a structural platform to probe the molecular mechanisms of AF-1 dependent PPARγ functions.

## RESULTS

### Structural comparison of the disordered AF-1 vs. structured LBD

PPARγ2 (referred to as PPARγ herein), the major isoform expressed in adipocytes ^26^, contains an N-terminal AF-1 domain also known as the A/B region encompassing residues 1-135 that is predicted to be structurally disordered (**Fig. 1A**). Experimental support for AF-1 structural disorder comes from high solvent D_2_O exchange observed by HDX-MS ^12^, which likely explains why crystal structures that include full-length PPARγ do not show AF-1 electron density (**Fig. 1B**). We validated these observations using several biophysical experimental approaches.

**Figure 1.**
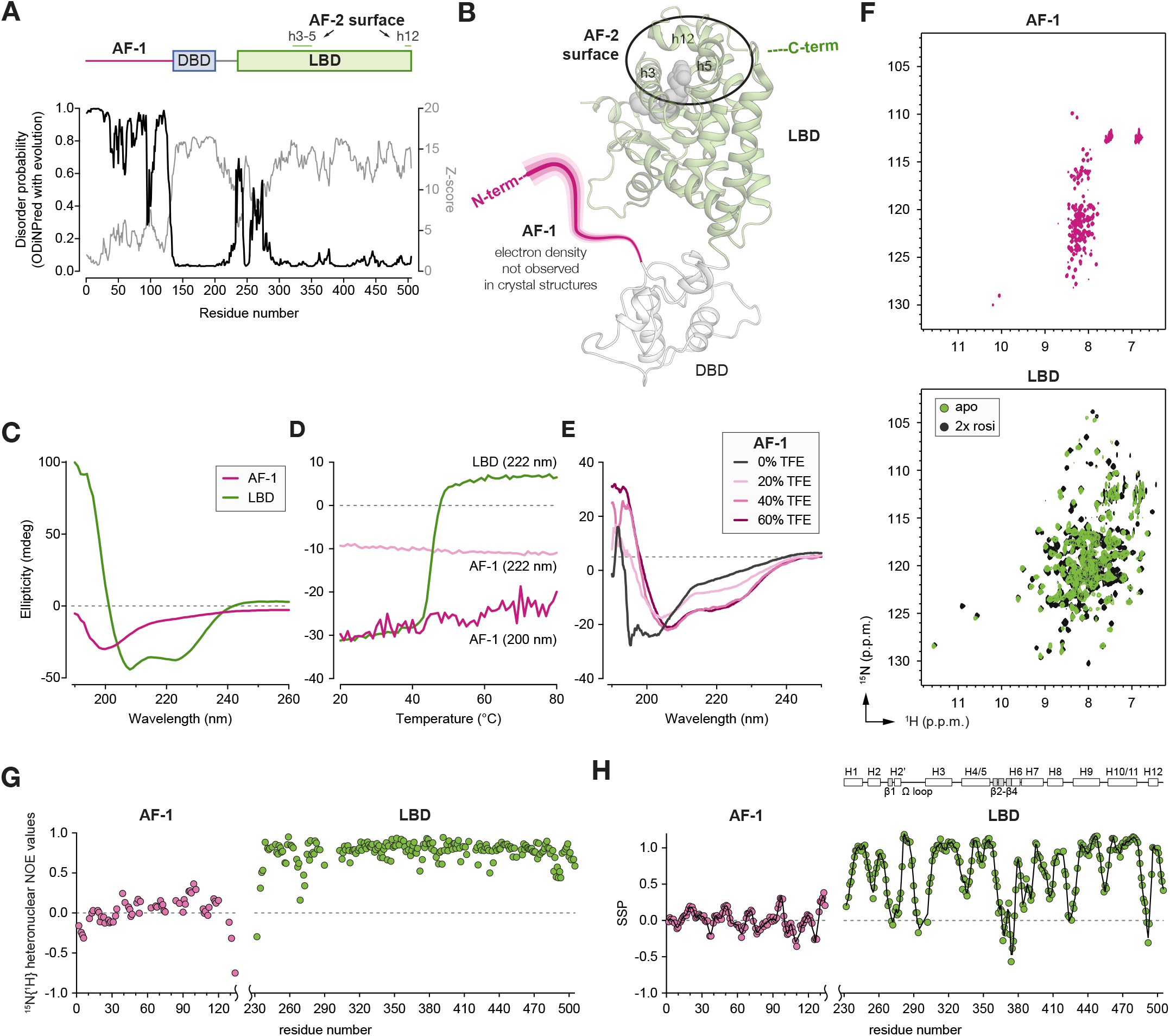
Biophysical characterization of the PPARγ AF-1 and LBD. (**A**) Domain architecture of human PPARγ isoform 2 (PPARγ2) and ODiNPred disorder prediction plot. (**B**) Crystal structure of full-length PPARγ (PDB 3DZY, chain D) highlighting the LBD (green) and the msising disordered AF-1 (pink). (**C**) Circular dichroism (CD) spectra of AF-1 and LBD. (**D**) Temperature-dependent CD thermal denaturation data of AF-1 and LBD. (**E**) CD spectra of the AF-1 collected with increasing concentrations of trilfluoroethanol (TFE). (**F**) 2D [^1^H,^15^N]-HSQC NMR spectrum of AF-1 (left) and 2D [^1^H,^15^N]-TROSY-HSQC NMR spectra of LBD in the apo and rosiglitazone-bound forms (right). (**G**) ^15^N{^1^H} heteronuclear NOE values of AF-1 and LBD. (**H**) Secondary structure propensity (SSP) values calculated from Cα and Cβ chemical shifts of the AF-1 and LBD.

Circular dichroism (CD) spectroscopy of the AF-1 reveals a random coil profile (**Fig. 1C**) with no temperature-dependent changes caused by domain unfolding, unlike the structured three-layer a-helical sandwich fold LBD that unfolds in a temperature-dependent manner (**Fig. 1D**). However, CD analysis of the AF-1 in the presence of increasing trifluoroethanol (TFE) shows an increase in a-helical signature (**Fig. 1E**), suggesting there may be transient secondary structure in the AF-1. 2D [^1^H,^15^N]-HSQC NMR analysis reveals poor amide spectral dispersion for the AF-1 characteristic of a disordered protein, whereas the ligand-free/ apo and ligand/agonist-bound LBD shows well dispersed amide chemical shifts (**Fig. 1F**). ^15^N{^1^H} heteronuclear NOE analysis, which reports on fast (picosecond time scale) backbone mobility ^27^, reveals low values for the AF-1 indicating large amplitude backbone motions compared to the LBD (**Fig. 1G**). Analysis of backbone Ca and Cp NMR chemical shifts, which predict secondary structure propensity in IDPs and folded proteins ^28^, reveals regions of the AF-1 that have low a-helical and p-sheet propensity compared, consistent with the TFE CD data, to the robust secondary structure present in the LBD (**Fig. 1H**). Taken together, these data indicate that although the AF-1 is structurally disordered, there are regions with transient or lowly populated secondary structure.

### Negative charge repulsion influences AF-1 compactness

Disordered proteins can adopt extended conformations with no intradomain contacts or compact conformational ensembles with transient or robust intradomain contacts, structural features that can be detected using paramagnetic relaxation enhancement (PRE) NMR methods ^29^. We used site directed mutagenesis to introduce a cysteine residue at several locations in the AF-1, which lacks native cysteine residues, and attached the cysteine-reactive nitroxide spin label MTSSL to each construct (D11C, S22C, D33C, D61C, A91C, S112C). We collected 2D [^1^H,^15^N]-HSQC NMR data in the paramagnetic and diamagnetic state and calculated peak intensity ratios (I_PRE_ = I_para_/I_dia_) to reveal AF-1 residues that are in close structural proximity to the MTSSL label (**Fig. 2A**). Generally, residues with I_PRE_ = 0 correspond to a distance <12Å, and I_PRE_ > 0 and < 1 correspond to a distance between 13-25Å where the peak intensity decrease is proportional to 1/r^6^ from the unpaired electron ^30^.

I_PRE_ values from AF-1 with an MTSSL spin label at D11 near the N-terminus (D11C-MTSSL) are consistent with an extended conformation with no significant long-range contacts as the profile is similar to a predicted I_PRE_ profile from an extended AF-1 ensemble calculated using *flexible-meccano* ^31^. In contrast, MTSSL placement at other regions within the AF-1 reveals long-range contacts given the experimental I_PRE_ values deviate from predicted extended I_PRE_ AF-1 profiles. Control experiments where ^15^N-labeled AF-1 is mixed with ^14^N/ MTSSL-labeled AF-1 show no PRE effects, indicating the PREs observed are due to intradomain AF-1 interactions and not interacts between distinct AF-1 molecules. Notably, the experimental PRE NMR profiles indicate the most robust intradomain interactions occur for residues where the MTSSL is placed between D33 and S112, suggesting most of the intradomain AF-1 interactions occur within this region.

**Figure 2.**
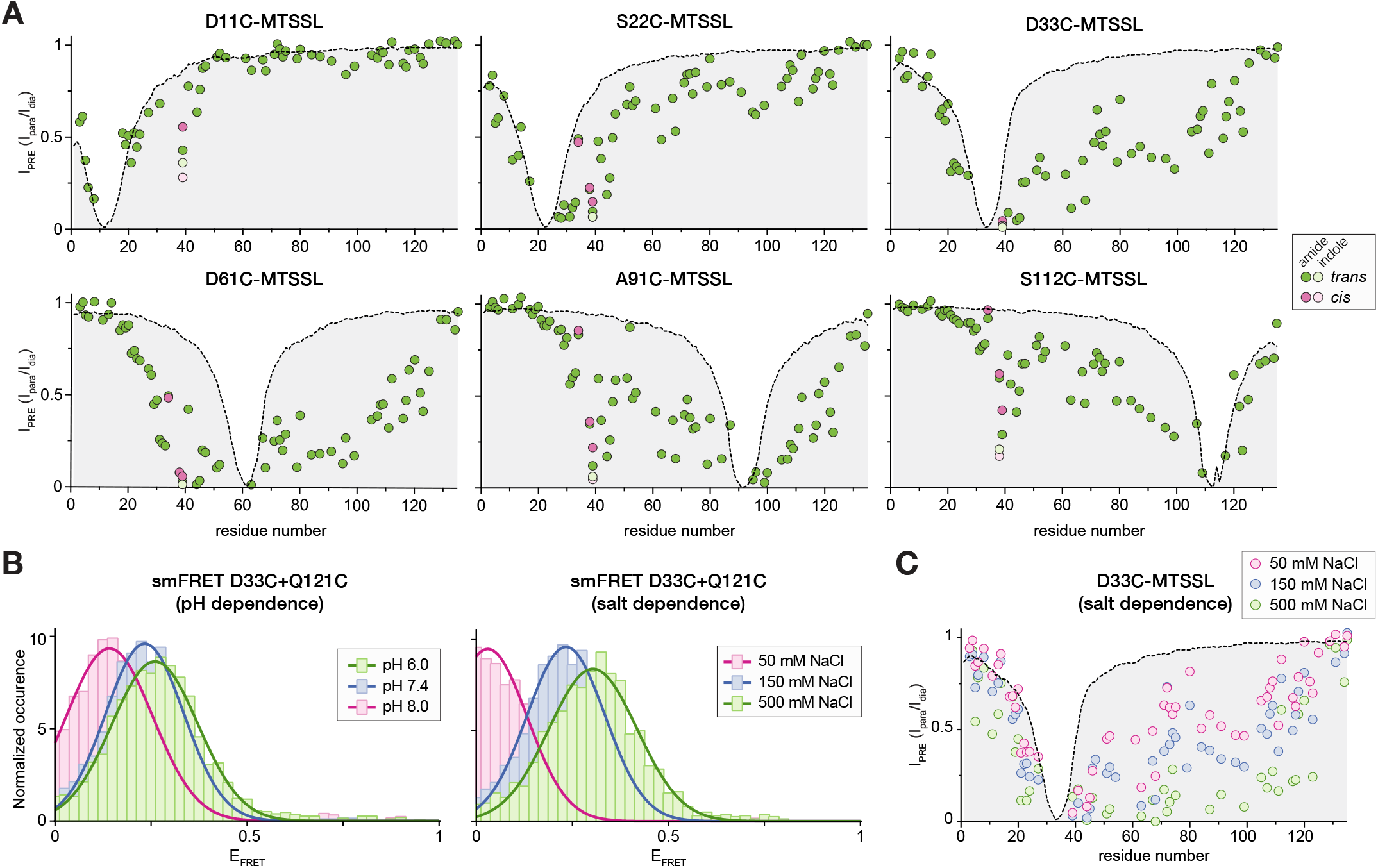
PRE NMR and smFRET analysis of the AF-1 structural ensemble. (**A**) Plots of experimental IPRE values (circles) calculated from peak intensities from 2D [^1^H,^15^N]-HSQC NMR spectra and calculated I_PRE_ profiles (dotted line with gray shaded area) from flexible-meccano simulations of extended AF-1 conformers for MTSSL-conjugated AF-1 constructs as noted. (**B**) Two-color smFRET data of D33C+Q121C double mutant AF-1 conjugated with Alexa Fluor 488 and 647 maleimide as a function of pH (left) or salt concentration (right). (**C**) Plot of experimental I_PRE_ values for the D33C-MTSSL AF-1 construct from 2D [^1^H,^15^N]-HSQC NMR spectra collected as a function of salt concentration (circles); calculated I_PRE_ profile for the extended D33C-MTSSL AF-1 conformers from (**A**) is also shown.

The sequence composition of the AF-1 shows a net negative charge (**Supplementary Fig. S1**) and a calculated pI of 4.17, which led us to hypothesize negative charge-charge interactions may influence the relative compactness of the AF-1 conformational ensemble. To test this, we performed smFRET using a two-color approach ^32,33^ with an AF-1 double mutant construct in which cysteines were placed near the N-terminus (D33C) and C-terminus (Q121C). Increasing FRET efficiency (E_FRET_) between the fluorophores attached to the D33C and Q121C sites occurs at lower pH or increasing salt concentration (**Fig. 2B**), indicating that salt neutralization of the negatively charged AF-1 side chains results in a more compact AF-1 conformation. Salt-dependent PRE NMR analysis of the AF-1 with the MTSSL spin label placed at D33C validated these findings since intramolecular PREs increased with increasing salt concentration (**Fig. 2C**). Taken together, the smFRET and PRE NMR data indicate the AF-1 transiently samples compact conformations that can be fine-tuned by changes in negative charge repulsion.

### AF-1 cis/trans isomerization via an evolutionarily conserved Trp-Pro motif

During backbone NMR chemical shift assignment of the AF-1, we noticed a region that displayed two populations of chemical shifts encompassing residues T34-N42 (^34^TEMPFWPTN^42^) — a sequence conserved among 350 PPARγ isoforms and orthologs from mammals, birds, and reptiles (**Fig. 3A**). The presence of two NMR-detected populations indicates this region slowly interconverts between at least two structural conformations on the millisecond-to-second time scale, a chemical exchange process that can be detected using ZZ-exchange NMR methods ^34^. Full-length PPARγ2 contains a single tryptophan residue (W39) that is present in the AF-1. However, two tryptophan side-chain indole peaks are observed in 2D [^1^H,^15^N]-HSQC NMR data of ^15^N-labeled AF-1 with populations of ∼85% and ∼15% (**Fig. 3B**). ZZ-exchange experiments at elevated temperatures revealed an increasing population of cross-correlated W39 indole peaks, indicating this region of the AF-1 slowly interconverts or isomerizes between two structural conformations on a timescale (τ_ex_) of milliseconds-to-seconds at temperatures above 35ºC and seconds-to-minutes below 35ºC.

**Figure 3.**
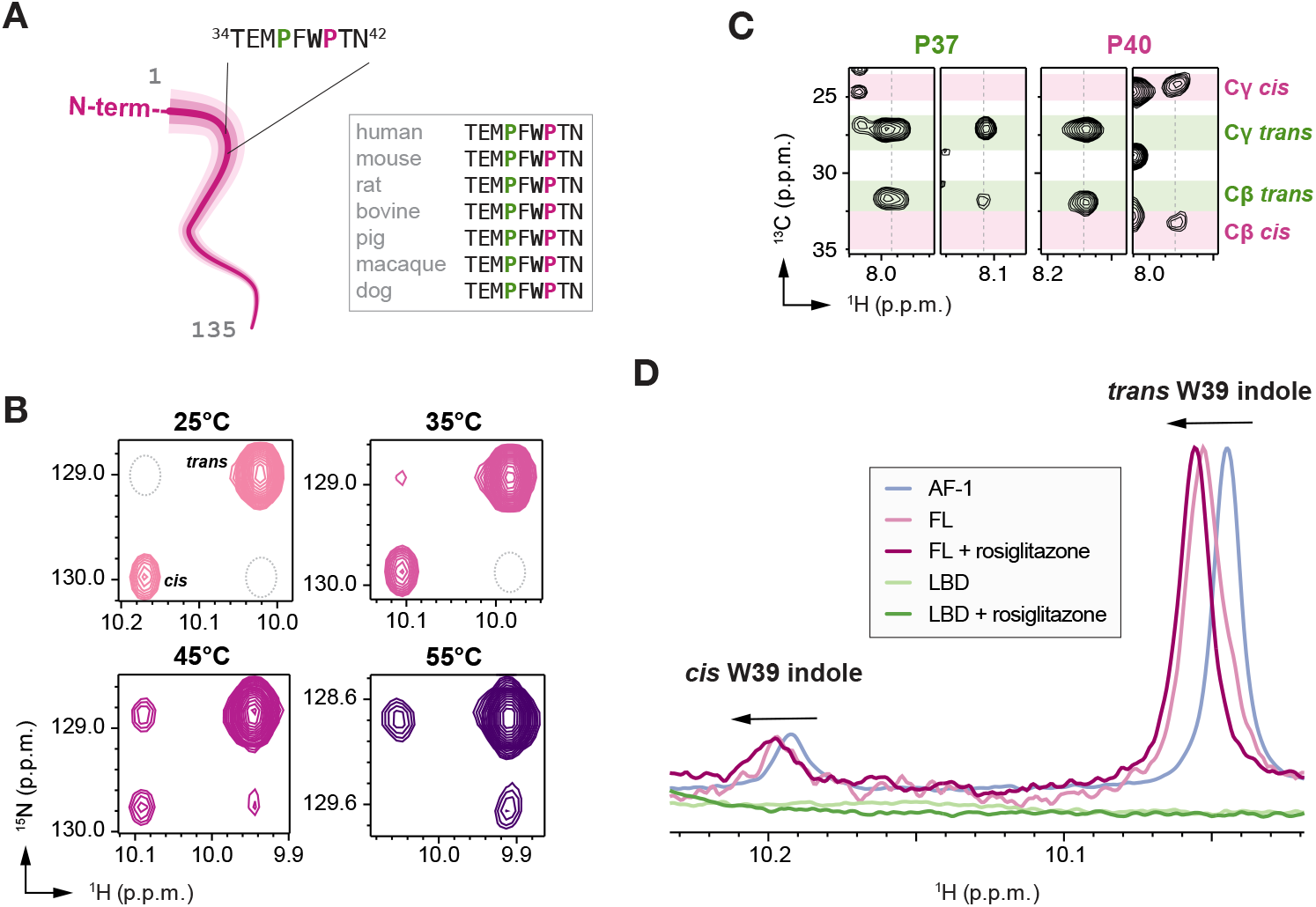
Proline cis/trans isomerization populates two long-lived AF-1 conformations. (**A**) Residues within the AF-1 displaying two populations of NMR peaks localize to a region containing two proline residues (P37, green; P40, pink), one of which is adjacent to a tryptophan residue (W39). (**B**) 2D [^1^H,^15^N]-ZZ-exchange NMR data (delay = 2 s) focused on the W39 indole peaks shows cross-correlated peaks appearing at elevated temperatures that are missing at lower temperatures (dashed gray oval). (**C**) Strips from 3D CC(CO)NH NMR data focused on P37 (i-1 to F38 amides) and P40 (i-1 to T41 amides). Average chemical shift ranges for *trans* and *cis* confomers of proline Cγ and Cβ atoms are displayed in green (*trans*) and pink (*cis*). (**D**) Overlay of 1D [^1^H]-NMR spectra of AF-1, LBD, or full-length (FL) PPARγ with or without addition of 2x rosiglitazone focused on the W39 indole peaks. Black arrows denote downfield shifting of the W39 indole NMR peak positions of FL vs. AF-1 samples.

Two proline residues (P37 and P40) located within this slowly isomerizing region led us to hypothesize that proline *cis/trans* isomerization contributes to the mechanism. CC(CO)NH NMR data, which provide chemical shifts proline Cβ and Cγnuclei—the values of which are predictive of *cis* or *trans* proline conformations ^35,36^ — revealed that the P40 backbone populates *cis* and *trans* conformations, whereas P37 populates only the *trans* conformation (**Fig. 3C**). These NMR data pinpoint the origin of the slowly isomerizing AF-1 conformational switch to the W39-P40 sequence, a dipeptide motif (Trp-Pro) previously shown to enrich the *cis* isomer ^37^.

To determine whether the two long-lived Trp-Prof conformations we observed in AF-1 protein occurs in full-length PPARγ, we compared 1D [^1^H]-NMR spectra of AF-1 and full-length PPARγ. Focusing on the indole peaks of W39 (**Fig. 3D**), two peaks are observed in full-length PPARγ that are shifted downfield (i.e., to the left) relative to the peaks observed in the AF-1 alone. Intriguingly, addition of the agonist ligand rosiglitazone, which stabilizes an active LBD conformation, shifts the full-length PPARγ W39 indole peaks further downfield. These data indicate that the presence of the LBD, as well as ligand binding to the LBD, affects the chemical environment of the AF-1, which we hypothesized may occur through an interdomain interaction between the AF-1 and LBD.

### AF-1/LBD interdomain interactions in full-length PPARγ

To extend the 1D NMR findings indicating the LBD influences the conformation of the AF-1, we used sortase A-mediated protein ligation and segmental isotope labeling ^38,39^ to generate full-length PPARγ protein where only the AF-1 is ^15^N-labeled and visible in 2D [^1^H,^15^N]-HSQC NMR data. Overlay of 2D NMR spectra of segmentally [^15^N-labeled AF-1] full-length PPARγ construct and the isolated ^15^N-labeled AF-1 reveals select chemical shift and line broadening changes (**Fig. 4A**). Notably, NMR samples of ^15^N-labeled AF-1 alone show a clear solution. However, the sortase-ligated full-length PPARγ NMR sample shows a condensed phase separated solution well dispersed throughout the entire NMR tube, consistent with published observations that the DBD mediates phase separation of PPARγ *in vitro* and in cells ^40^. We therefore performed NMR studies using sortase-ligated full-length PPARγ in addition to NMR titration studies using isolated AF-1 and LBD proteins (*vide infra*), which do not phase separate due to the absence of the DBD, to isolate effects arising from potential interactions between proteins in the phase separated state vs. interdomain effects in full-length PPARγ such as a physical interaction between domains or conformational change.

**Figure 4.**
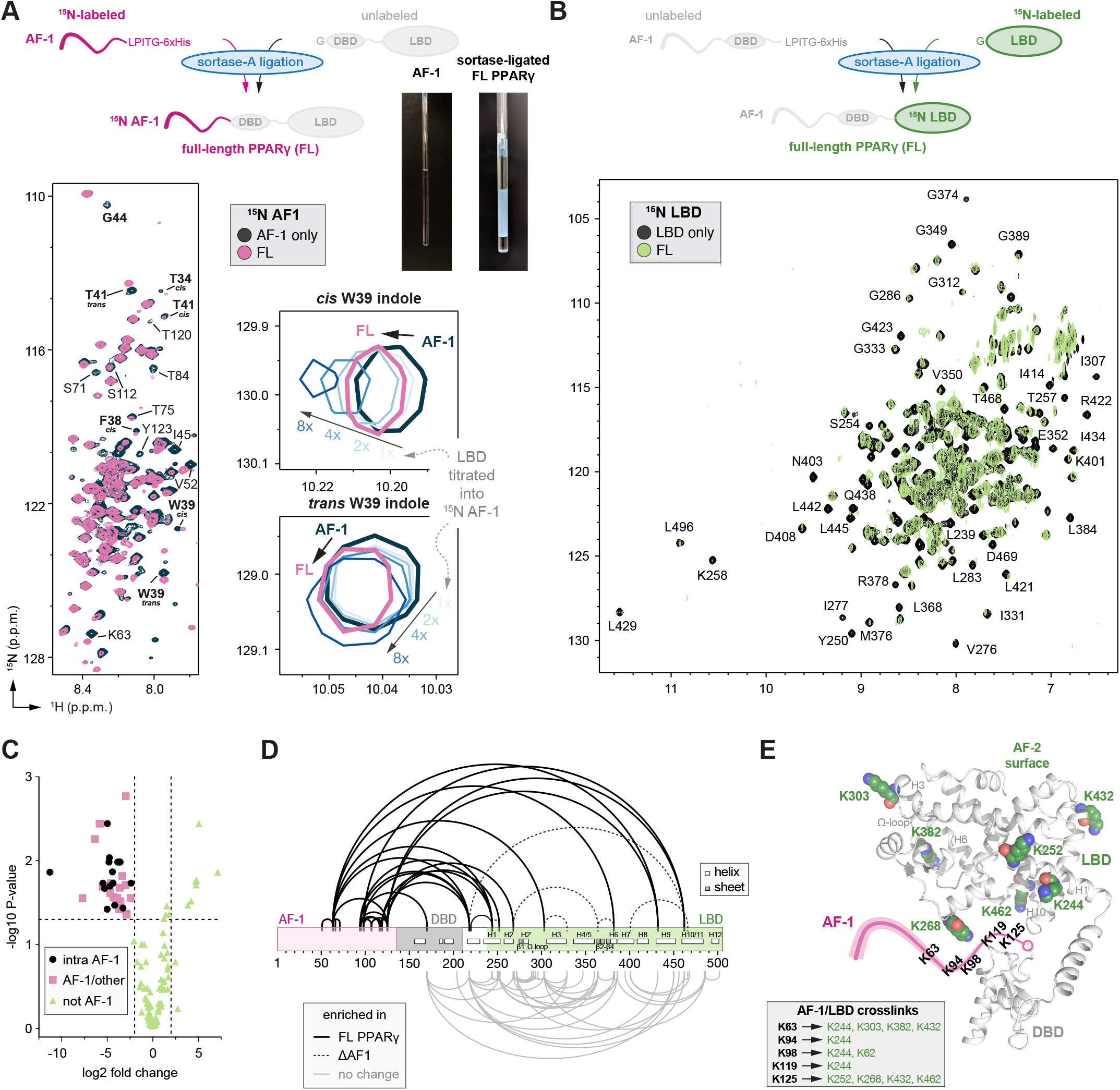
NMR and XL-MS reveal the AF-1 and LBD interact in full-length PPARγ. (**A**) Overlay of 2D [^1^H,^15^N]-HSQC NMR spectra comparing ^15^N AF-1 to sortase A-ligated full-length PPARγ where only the AF-1 (residues 1-135) is ^15^N-labeled. Residues with notable differences are annotated, and residues near the Trp-Pro (W39-P40) motif are in bold font. (**B**) Overlay of 2D [^1^H,^15^N]-TROSY-HSQC NMR spectra comparing ^15^N LBD to sortase A-ligated full-length PPARγ in rosiglitazone-bound forms (2x) where only the LBD (residues 231-505) is ^15^N-labeled. Residues with notable differences are annotated. (**C**) Plot of the P-value vs. fold change of DSSO crosslinks from the differential XL-MS analysis. (**D**) DSSO crosslinks detected in differential XL-MS analysis of full-length PPARγ vs. AF-1 truncation (ΔAF-1). (**E**) LBD residues involved in AF-1 crosslinks plotted on the AlphaFold2 structure of PPARγ (residues 136-505 shown) with a schmatic of the AF-1, which is missing in crystal structures of full-length PPARγ, shown in pink.

Similar to the 1D NMR analysis, the two W39 indole peaks are shifted downfield (i.e., to the left) in 2D [^1^H,^15^N]-HSQC NMR data of sortase-ligated full-length PPARγ containing an ^15^N-labeled AF-1 relative to isolated ^15^N-labeled AF-1 (). Moreover, titration of unlabeled LBD into ^15^N-labeled AF-1 showed a similar chemical shift perturbation (CSP) pattern for the W39 indole peaks, which not only confirms a direct AF-1/LBD interaction but also suggests the interaction between isolated domains is similar to the interaction in full-length PPARγ. Other residues within and near near the slowly exchanging Trp-Pro motif also show CSPs including well resolved backbone amide peaks corresponding to T34 (*cis*), F38 (*cis*), W39 (*cis* and *trans*), T41 (*cis* and *trans*), and G44. Furthermore, other residues within regions involved in AF-1 intramolecular PRE contacts show chemical shift and peak line broadening changes.

We also collected 2D [^1^H,^15^N]-TROSY-HSQC NMR data of sortase ligated full-length PPARγ containing an ^15^N-labeled LBD relative to isolated ^15^N-labeled LBD in the rosiglitazone-bound states (**Fig. 4B**). NMR peak line broadening is observed for residues comprising several distinct LBD surfaces including the AF-2 coregulator interaction surface (L345, K347, Y348, G349, G423, R425, L429, L496, L497, K502) and the β -sheet region (F275, I277, G374, M376). These line broadening effects could be due to a combination of the larger molecular weight of the isolated LBD vs. segmentally ^15^N-labeled LBD in full-length PPARγ and possible interdomain interactions that occur within the context of full-length PPARγ. Furthermore, some peaks show notable CSPs including F275, G389, K401, G427, L428, and H453, which includes the AF-2 surface; these residues experience a different chemical environment relative to the isolated LBD.

We performed chemical crosslinking mass spectrometry (XL-MS) as an orthogonal method to structurally map the interdomain interactions in full-length PPARγ. We compared DSSO-crosslinked samples of full-length PPARγ to a truncated construct without the AF-1 (ΔAF-1) containing the DBD-hinge-LBD to delineate crosslinks enriched or decreased in the presence or absence of the AF-1 (**Fig. 4C**). Thirty-eight crosslinks were enriched in full-length PPARγ, including sixteen unique intradomain AF-1 crosslinks and thirteen unique AF-1/LBD crosslinks (**Fig. 4D**). Crosslinks to the AF-1 localize primarily to the C-terminal half of the AF-1 between K94 and K125, which contains five out of six lysine residues present in the AF-1. Seven LBD residues involved in interdomain AF-1 crosslinks include residues near the p-sheet surface on helix 1 (K244, K252), helix 2 (K268), Ω -loop (K303), helix 6 (K382), and helix 10 (K462), and near the AF-2 surface (K432). These LBD crosslinks are consistent with the putative location of the AF-1 region in the crystal structure of full-length PPARγ where electron density for the AF-1 was not observed (**Fig. 4E**) ^12^. Six interdomain DBD crosslinks were detected, including one AF-1/DBD crosslink and five DBD/LBD crosslinks. Furthermore, seven crosslinks were enriched in ΔAF-1 PPARγ, six of which are LBD/LBD near the p-sheet surface. These XL-MS data indicate that the AF-1, and by corollary its removal, structurally affects the LBD conformation.

### AF-1 Trp-Pro motif is a major determinant in LBD interaction

Although the DSSO-crosslinked XL-MS data reveal interdomain interactions between the C-terminal half of the AF-1 and LBD, the differential AF-1 vs. sortase-ligated full-length NMR data suggest a more extensive interaction involving other regions of the AF-1 that lack lysine residues, in particular residues near the Trp-Pro motif. To better characterize the AF-1/LBD interaction, we performed NMR chemical shift structural footprinting analysis using isolated AF-1 and LBD, which also allowed us to remove phase separation contributions to the NMR spectra that occur in studies of full-length PPARγ when the DBD is present.

To map the LBD-binding interface within the AF-1, we collected 2D [^1^H,^15^N]-HSQC NMR data (**Fig. 5A**) and 2D [^13^C,^15^N]-(HACA)CON NMR data (**Fig. 5B**) using ^15^N- and ^13^C,^15^N-labeled AF-1, respectively, in the absence and presence of LBD. Addition of LBD caused CSPs (fast exchange on the NMR time scale) and peak line broadening (intermediate exchange on the NMR time scale) for select residues, confirming a direct interaction between the isolated domains. Three notable features are apparent in these NMR structural footprinting data. First, AF-1 residues most affected displaying the largest CSPs and peak line broadening changes include F38, W39, P40, and T41, pinpointing the region containing the slowly exchanging Trp-Pro motif as a major determinant in LBD interaction. Similar to the 1D and 2D differential NMR data of full-length PPARγ and sortase-ligated full-length PPARγ, the W39 indole side chain peaks are shifted downfield with increasing LBD concentrations. These observations indicate the interaction between the isolated AF-1 and LBD interaction is similar to what occurs within the context of full-length PPARγ even in the context of phase separation. Second, AF-1 residues distal in primary sequence from, but involved in intramolecular PRE contacts to, the Trp-Pro region show CSPs and peak line broadening. These observations suggest a more extensive LBD interaction interface or allosteric conformational changes that occur in the AF-1 upon LBD interaction. Third, increasing LBD concentrations primarily cause CSPs on the fast exchange NMR time scale (peak shifting) for the Trp-Pro region *trans* conformation peaks, whereas the *cis* conformation peaks display a combination of fast and intermediate exchange (peak disappearance). This difference in exchange on the NMR time scale suggests that the *cis* AF-1 conformation may interact with the LBD with higher affinity and/or different interaction kinetics than the *trans* AF-1 conformation.

**Figure 5.**
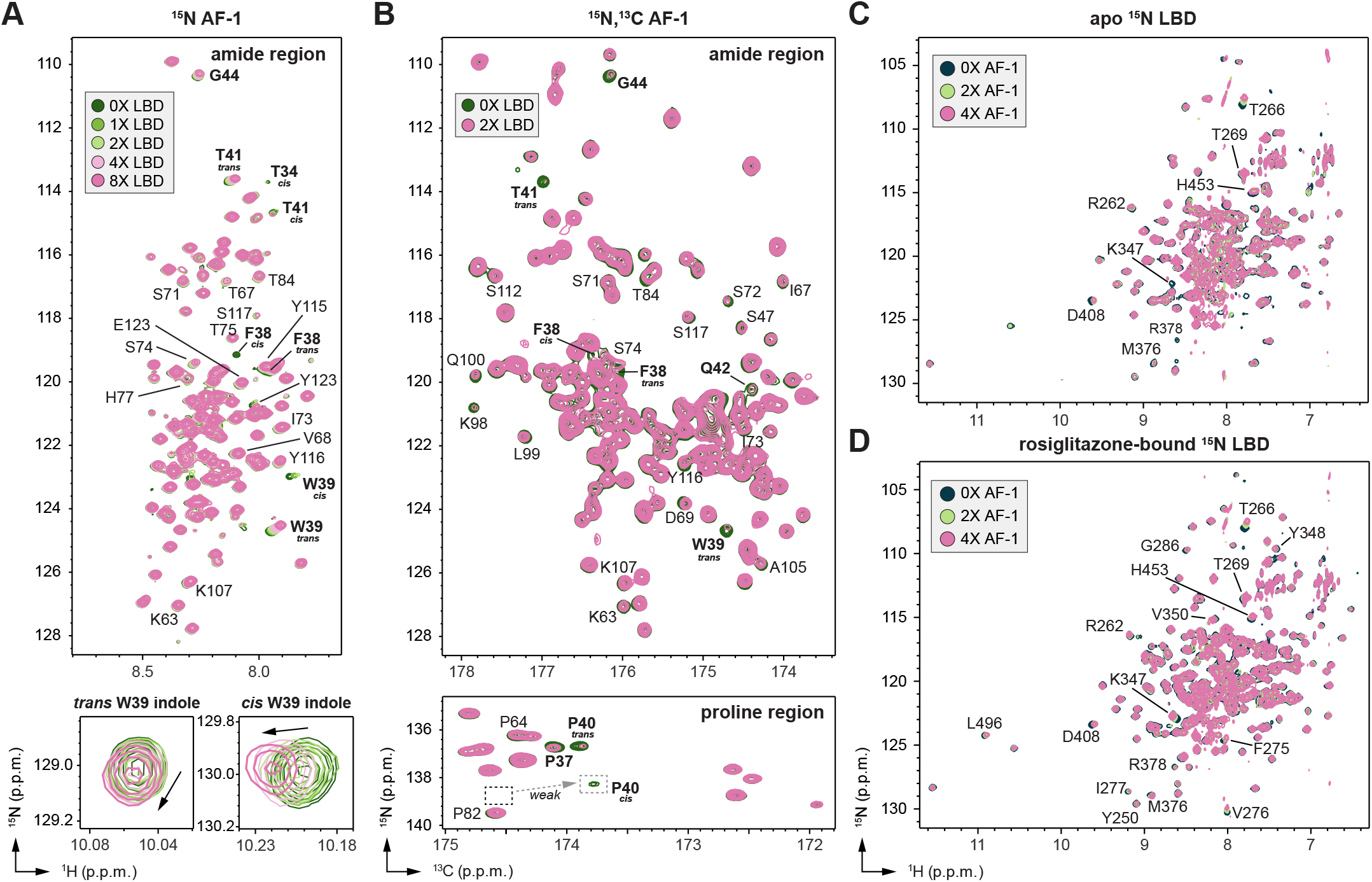
NMR structural footprinting of AF-1/LBD interaction using isolated domains. (**A**) Overlay of 2D [^1^H,^15^N]-HSQC NMR spectra of ^15^N-labeled AF-1 with increasing concentrations of LBD. (**B**) Overlay of 2D [^13^C,^15^N]-(HACA)CON NMR spectra of ^13^C,^15^N-labeled AF-1 containing an N-terminal 6xHis+TEV tag with or without LBD. Residues with notable differences are annotated, and residues near the Trp-Pro (W39-P40) motif are in bold font. (**C**,**D**) Overlay of 2D [^1^H,^15^N]-TROSY-HSQC NMR spectra of (**C**) ^15^N-labeled apo-LBD or (**D**) ^15^N-labeled rosiglitazone-bound LBD with increasing concentrations of AF-1. Residues with notable differences are annotated, and residues near the Trp-Pro (W39-P40) motif are in bold font.

### Trp-Pro AF-1 motif interacts with two LBD surfaces

To map the AF-1-binding interface within the LBD, we collected 2D [^1^H,^15^N]-TROSY-HSQC NMR data of ^15^N-labeled PPARγ LBD in the absence and presence of AF-1. Titration of AF-1 into ^15^N-labeled apo-LBD revealed chemical shift perturbations for several NMR peaks (**Fig. 5C**). Residues that show the largest chemical shift changes colocalize near the p-sheet surface and include R262, T266, and T269 on helix 2; D408 on the helix 7-8 loop; and H453 on the helix 9-10 loop. However, apo/ligand-free PPARγ LBD dynamically exchanges between transcriptionally active and repressive conformations on the intermediate exchange NMR time scale resulting in missing or very broad peaks for about half of the LBD including residues within and near the orthosteric ligand-binding pocket and AF-2 surface ^6,41^ due to dynamic exchange between active and repressive LBD conformations ^7,8^. Binding of a full agonist such as rosiglitazone stabilizes an active LBD conformation where nearly all NMR peaks are visible ^5,6,8^. TItration of AF-1 into rosiglitazone-bound^15^N-labeled LBD revealed additional chemical shift perturbations in regions that are stabilized in the active conformation (**Fig. 5D**), including residues within the p-sheet surface (F275, V276, I277, G286, R378) and AF-2 coregulator interaction surface (K347, Y348, and V350 on helix 5; and L496 on helix 12).

Because NMR chemical shift footprinting analysis reports on residues directly involved in the AF-1/LBD interaction as well as allosteric conformational changes that occur upon binding, we used intermolecular PRE NMR to specifically map where the AF-1 region containing the Trp-Pro motif interacts on the LBD. We collected 2D [^1^H,^15^N]-TROSY-HSQC PRE NMR data of rosiglitazone-bound ^15^N-labeled LBD in the presence of D33C-MTSSL AF-1 (**Fig. 6A**), which places the MTSSL group near the Trp-Pro motif. LBD residues with decreased I_PRE_ values (**Fig. 6B**) cluster at two distinct surfaces (**Fig. 6C**): a large group at the AF-2 coregulator interaction surface, and a smaller group near the β -sheet surface.

**Figure 6.**
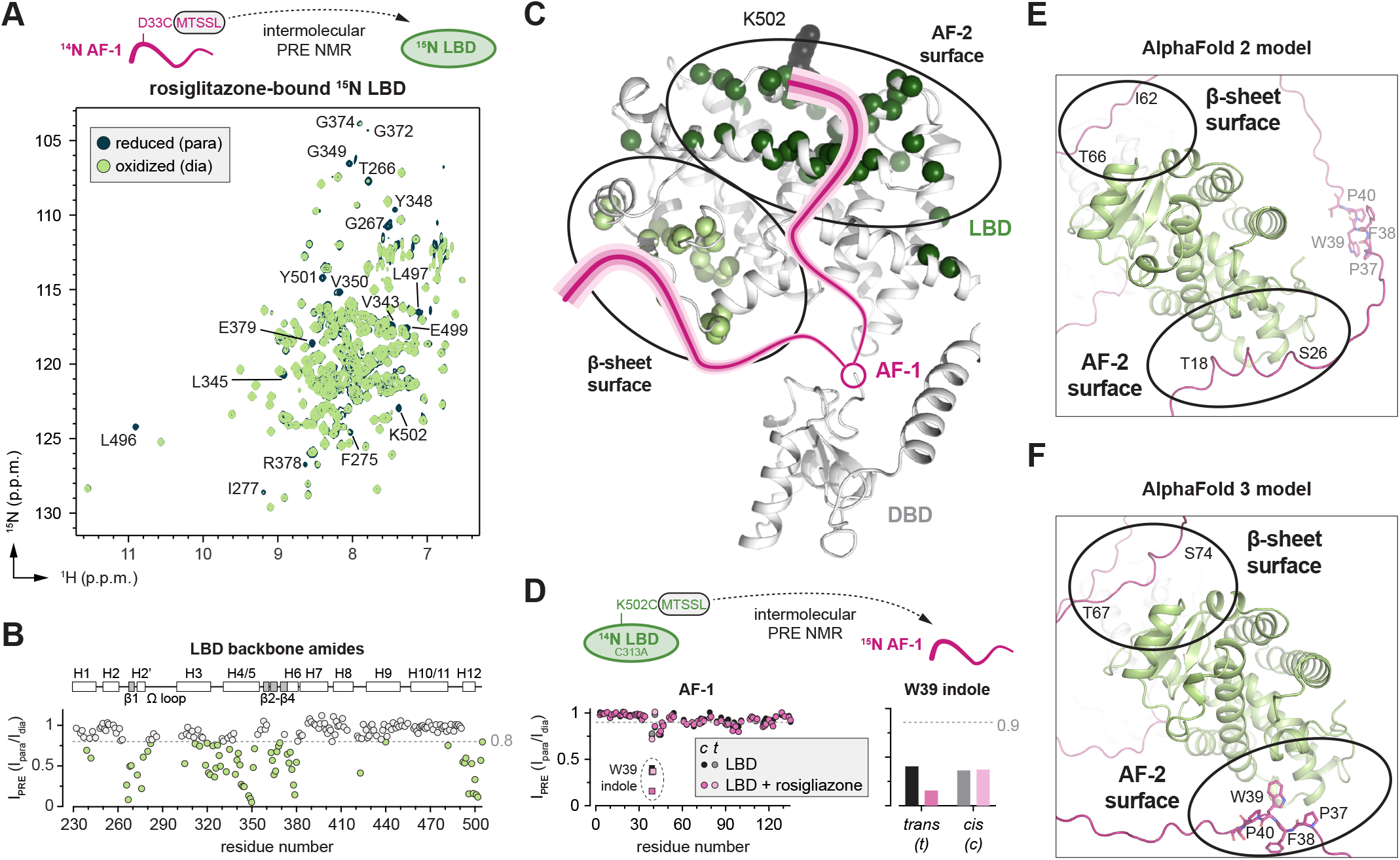
PRE NMR analysis reveals the AF-1 Trp-Pro motif interacts with two LBD surfaces. (**A**) Overlay of 2D [^1^H,^15^N]-TROSY-HSQC PRE NMR spectra of ^15^N-labeled rosiglitazone-bound LBD with 2x D33C-MTSL AF-1 in the paramagnetic (oxidized) and diamagnetic (reduced) states. Residues with notable differences are annotated. (**B**) Plot of I_PRE_ values calculated from the PRE NMR spectra in (**A**) vs. residue number. Secondary structural elements of the LBD are shown above the plot. (**C**) LBD residues with I_PRE_ values ≤ 0.8 in (**B**) displayed on the AlphaFold2 structure of PPARγ (residues 136-505) map to two distinct surfaces (light vs. dark spheres; black circles); the putative location of the AF-1, which is missing in crystal structures of full-length PPARγ, is shown in pink. (**D**) Plot of backbone amide and W39 indole I_PRE_ values calculated from the PRE NMR spectra of ^15^N-labeled AF1 with 2x K502C-MTSSL LBD vs. residue number. (**E**,**F**) Models of full-length PPARγ2 predicted by AlphaFold 2 (**E**) and AlphaFold 3 (**F**).

To confirm the AF-2 surface interaction with the Trp-Pro motif, we performed an analagous intermolecular PRE NMR experiment using ^15^N-labeled AF-1 in the presence of an MTSSL-labeled double mutant LBD (C313A/K502C) we previously generated ^8,42,43^. The C313/K502C mutant LBD (1) contains a cysteine residue within helix 12 (K502C) that is solvent exposed in the active AF-2 surface conformation when bound to rosiglitazone and (2) removes the only cysteine residue in the LBD (C313A) to enable site-specific covalent labeling to the K502C on AF-2 helix 12. AF-1 residues with decreased I_PRE_ values (**Fig. 6D**) cluster primarily within and near the Trp-Pro motif, with additional albeit smaller PRE effects in regions involved in intradomain AF-1 contacts detected by PRE NMR (**Fig. 2A**). Notably, addition of rosiglitazone resulted in a lower I_PRE_ value for the W39 indole *trans* conformation compared to the *cis* conformation, which could indicate an interaction preference with the Trp-Pro motif *trans* AF-1 conformation to an agonist-stabilized active LBD conformation.

Taken together, the intermolecular PRE NMR data corroborate and extend the NMR structural footprinting data and XL-MS data, which show that the p-sheet and AF-2 LBD surfaces are involved in the interdomain AF-1 interaction. Furthermore, our studies pinpoint the slowly exchanging AF-1 Trp-Pro motif, which populates two long-lived AF-1 structural conformations, as a major interaction determinant with the AF-2 coregulator interaction surface in the LBD. AlphaFold 2 ^18^ prediction of full-length PPARγ2 (**Fig. 6E**), suggest an AF-1 interaction with the LBD/AF-2 surface but via an AF-1 region not supported by our NMR data encompassing residues 18-25, which is present in the PPARγ2 isoform but not the shorter PPARγ1 isoform. Curiously, however, AlphaFold 3 ^44^ predicts (**Fig. 6F**) an interaction between the AF-1 Trp-Pro motif (PFWP-containing region) and the LBD/AF-2 surface where the W39 side chain points into the AF-2 surface, consistent with our CSP AND PRE NMR data that show the largest CSPs occurring for the W39 backbone and indole amides. Furthermore, AlphaFold 2 and 3 predictions suggest an interaction between AF-1 residues in the 60s and 70s to the p-sheet surface of the LBD, of which the AlphaFold 3 predictions are better supported by NMR data where titration of LBD into 15N-labeled AF-1 causes more notable CSP for residues 70-75.

### Mutation of the AF-1 Trp-Pro motif impacts LBD interaction and transcription

We generated several mutant AF-1 constructs including single site W39A or P40A mutants and a quadruple P37A/F38A/W39A/P40A mutant (PFWP-to-AAAA) and used 2D [^1^H,^15^N]-HSQC NMR data to determine the relative *cis* and *trans* conformations of the Trp-Pro region (**Supplementary Fig. S2**). The single site mutations, W39A and to a larger degree P40A, suppress but do not completely abolish *cis/trans* isomerization of the Trp-Pro region (**Supplementary Table S1**). However, the quadruple PFWP-to-AAAA AF-1 mutant, which encompasses residues in the AF-1 region most affected in the LBD NMR CSP data, displays no detectable *cis* population for residues within the Trp-Pro region, indicating this mutant may provide a means to determine if there is a functional role of the Trp-Pro motif.

We collected 2D [^1^H,^15^N]-HSQC NMR data of ^15^N-labeled AF-1 PFWP-to-AAAA mutant in the absence and presence of increasing concentrations of LBD (**Fig. 7A**), which revealed CSPs for select residues distal from the Trp-Pro motif that were also affected when LBD is titrated into ^15^N-labeled AF-1 wild-type. However, the LBD does not appear to interact robustly with the mutated PFWP-to-AAAA region, as several new unassigned NMR peaks resulting from the alanine mutations show no significant or relatively minor CSPs. To determine if the Trp-Pro region influences PPARγ-mediated transcription, we performed a cellular transcriptional reporter assay where HEK293T cells were transfected with a luciferase reporter plasmid containing an idealized PPAR-binding DNA response element sequence (PPRE) along with expression plasmids for wild-type PPARγ or a full-length PPARγ mutant variant containing the PFWP-to-AAAA mutations (**Fig. 7B**). Transfection of wild-type PPARγ resulted in an increase in transcription, which was further increased upon addition of the PPARγ agonist ligand rosiglitazone. However, the PFWP-to-AAAA mutant PPARγ showed lower basal activity and diminished activation by rosiglitazone. These findings suggest that the Trp-Pro region plays an important role in the interdomain AF-1/LBD interaction and transcriptional activation mechanism of PPARγ.

**Figure 7.**
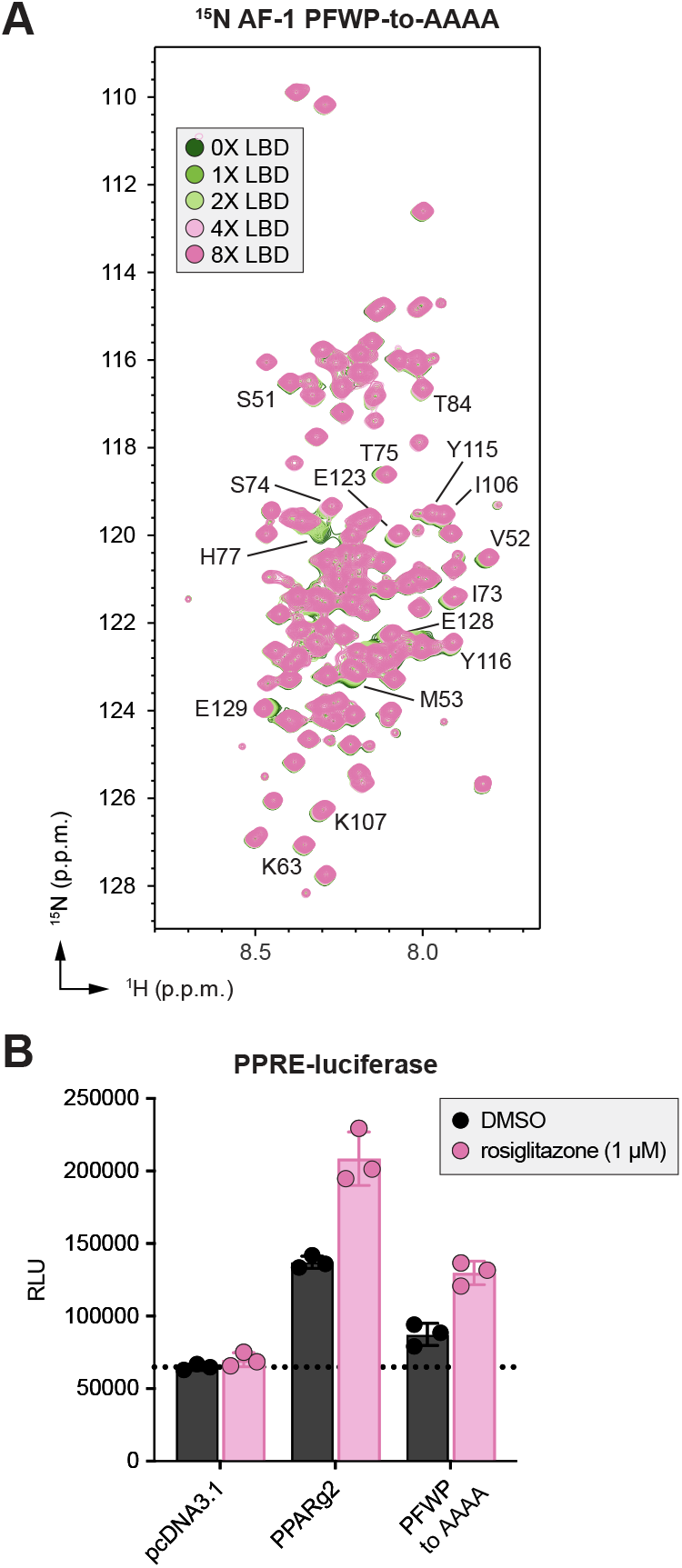
Mutation of the AF-1 Trp-Pro motif impacts LBD surfaces interaction and PPARγ-mediated transcription. (**A**) Overlay of 2D [^1^H,^15^N]-HSQC NMR spectra of ^15^N-labeled AF-1 PFWP-to-AAAA mutant with increasing concentrations of LBD. (**B**) Cell-based luciferase reporter assay in HEK293T cells transfected with full-length PPARγ or PFWP-to-AAAA mutant PPARg along with a PPRE-luciferase reporter plasmid treated with rosiglitazone (1 µM) or DMSO control (n=3). Bars and error bars represent the mean and s.d.; data representative of >2 experiments.

## DISCUSSION

The disordered N-terminal PPARγ AF-1 domain is an important regulator of PPARγ function. Previous studies have indicated the AF-1 negatively regulates PPARγ transcription and PPARγ-mediated adipogenesis via an AF-1/LBD interdomain functional communication mechanism ^20–24^. However, the structural basis for AF-1 activity, including its effects on the LBD, has remained elusive due to its disordered properties that make it difficult to study by X-ray crystallography or cryo-EM. Here, we used a combination of solution-based techniques to characterize the structural properties of the PPARγ AF-1. We found that although predominantly disordered, the AF-1 is characterized by transient long-range intramolecular contacts that can be enhanced under conditions that decrease negative charge repulsion. Thus, a change in AF-1 structure and compaction could occur under cellular states that would affect protein net charge, such as the presence of binding partners including other proteins, nucleic acids (mRNA and chromatin), or small molecule metabolites ^1^; upon post-translational modification of PPARγ ^45^; or cellular compartments with localized changes in ionic strength or pH such as phase separated biomolecular condensates ^40^. Long range interactions were also observed in structural studies of estrogen receptor alpha (ERa) AF-1 ^46,47^, which are modulated by phosphorylation ^48^. Thus, it will be interesting in future studies to determine how phosphorylation of the PPARγ AF-1 influences intradomain and interdomain interactions.

Our NMR analysis also revealed that the AF-1 contains an evolutionarily conserved Trp-Pro motif that undergoes *cis/trans* isomerization on a slow timescale (milliseconds to seconds) populating two long-lived AF-1 conformations. Proline *cis/trans* isomerization plays important roles in protein folding ^49^ and functions including isomerization-regulated ion channel pore opening ^50^, control of interdomain autoinhibitory interactions ^51–53^, and regulation of protein-protein interactions ^54,55^. Our discovery of the PPARγ AF-1 Trp-Pro motif interaction with the AF-2/ LBD is, to our knowledge, a unique example of a disordered domain undergoing *cis/trans* isomerization that participates in an interdomain interaction with a structured domain. The observation that the PFWP-to-AAAA mutation decreases basal and agonist-induced transcriptional activity of PPARγ and is involved in an AF-1/LBD interaction indicates isomerization of this region may have important cellular functions that warrants further study. Furthermore, our NMR data indicate that the LBD interaction with the *cis* and *trans* AF-1 PFWP motif conformations may occur with different kinetic exchange properties, and that the LBD may preferentially interact with the *trans* isomer in the presence of activating ligand. Thus, changes to the isomerization rate or relative isomer populations (i.e., by a proline isomerase enzyme) could also modulate the AF-1/LBD interaction. We recently found that the prolyl isomerase Pin1, which interacts with the PPARγ AF-1 ^56^, affects *cis/trans* isomerization of the Trp-Pro motif via a tethering mechanism ^57^, which could provide a mechanism to influence AF-1/LBD interactions, a future direction worth pursuing.

Published studies showed that the AF-1 inhibits ligand-dependent PPARγ activities ^20–24^ but did not determine whether the effects were due to a direct AF-1/LBD interaction. Notably, AF-1/NTD interactions with nuclear receptor LBDs have been reported or suggested for at least four other nuclear receptors including androgen receptor (AR) ^58–65^, ER ^66,67^, glucocorticoid receptor (GR) ^68^, and mineralocorticoid receptor (MR) ^69,70^. Our CSP and PRE NMR revealed that the AF-1 contacts the LBD at two distinct surfaces. Unlike the AF-1/AF-2 interaction identified for androgen receptor, in which an FXXLF motif within the AF-1 folds upon binding, the “fuzzy” interaction observed for PPARγ aligns with others previously identified for disordered proteins ^71,72^. Our results indicate the LBD AF-2 surface predominantly interacts with the region encompassing the AF-1 Trp-Pro motif, which is present in both major PPARγ isoforms, the longer adipocyte-specific γ2 isoform and the 28-residue shorter γ1 isoform that displays broader cell/tissue expression. Although our NMR data suggest the interaction affinity is weak when the isolated AF-1 and LBD are titrated together, which causes CSPs that occur on the fast-to-intermediate NMR time scale, within the context of full-length PPARγ the interaction is likely more robust since the two domains are physically tethered together. AF-1 interaction with the AF-2 surface could directly compete for coactivator binding, as previously observed for the AR and ER AF-1 ^65,67^, which could explain why AF-1 deletion increases the expression of PPARγ-regulated gene expression in adipocytes. Likewise, AF-1 interactions at the LBD p-sheet surface could interfere with ligand exchange into the orthosteric ligand-binding pocket ^4^, conformational changes associated with ligand binding, or interaction with other nuclear receptors ^12^ or other proteins. Further studies are warranted to explore the mechanistic basis by which the AF-1/LBD interaction affects these and other LBD-focused functions.

In addition to characterizing a direct interdomain interaction between the PPARγ AF-1 and LBD, our solution-based structural approach illuminates a platform to interrogate the activities of disordered nuclear receptor domains with extension to full-length proteins. While nearly all of the 48 human nuclear receptors contain unstructured AF-1 regions with important documented functional roles, the structural basis for their activities remain largely unexplored. Thus, it will be important to apply structural approaches such as those outlined here to gain a more comprehensive understanding of nuclear receptor activity.

## MATERIALS AND METHODS

### Plasmids and reagents

All plasmids and constructs use PPARγ 2 isoform numbering. For bacterial expression of proteins, DNA sequences encoding PPARγ AF-1 domain (residues 1-135), PPARγ LBD (residues 231-505), full-length PPARγ (residues 1-505), and PPARγ ΔAF-1 (residues 136-505) were inserted into pET45 or pET46 plasmids as tobacco etch virus (TEV) protease-cleavable N-terminal hexahistidine (6xHis) tag fusion proteins. Plasmids for in vitro sortase A-mediated protein ligation were generated using a published protocol ^38^; the N-terminal sortase A construct contained the AF-1 (residues 1-135) along with a 5-residue extension containing the sortase A consensus sequence (LPITG) followed by a 6xHis-tag; the C-terminal construct contained a TEV-cleavable N-terminal 6xHistag followed by the PPARγ DBD-hinge-LBD (residues 136-505), which upon TEV cleavage leaves an N-terminal Gly residue required for sortase A ligation. For mammalian cellular studies, plasmids including full-length PPARγ isoform 2 (residues 1-505) and pGL3-synthetic-PPRE-Luc2 luciferase reporter were previously reported ^73,74^. Mutant constructs were generated using site directed mutagenesis using PfuUltra II high fidelity polymerase (Agilent) and following manufacturer’s protocol. All plasmids were confirmed by Sanger sequencing (Genewiz) prior to use.

### Protein expression and purification

Proteins were expressed in *Escherichia coli* BL21(DE3) cells (Life Technologies). Full-length and ΔAF-1 PPARγ proteins were expressed using an autoinduction procedure where cells were grown for 6 hrs at 37 °C followed by addition of 0.1 mM zinc chloride, temperature reduction to 22 °C for 16 hrs. PPARγ AF-1, LBD, and sortase A-compatible proteins were either expressed in Terrific Broth (TB) media or M9 media supplemented with ^13^C-glucose and/or ^15^NH_4_Cl (Cambridge Isotope Labs, Inc.) followed by protein induction with 0.5 mM isopropyl ß-D-1-thiogalactopyranoside (Gold Biotechnology) at 18°C for 16 hrs. Cells were harvested by centrifugation, washed with phosphate buffered saline (PBS) buffer, and resuspended in a cell lysis buffer containing 40 mM potassium phosphate (pH 7.4), 500 mM potassium chloride, 15 mM imidazole, 0.5mM ethylenediaminetetraacetic acid (EDTA), 1 mM phenylmethylsulfonyl fluoride (PMSF), and Pierce protease inhibitor tablets (Thermo Scientific). Cells were lysed by sonication and the lysate was clarified by centrifugation at 14000 rpm for 45 min and filtered with a 0.2µm filter prior to loading into the Ni-NTA column. The protein was eluted against a 500 mM imidazole gradient through a Ni-NTA column, followed by overnight dialysis against a buffer without imidazole for TEV protease His tag cleavage at 4°C. The next morning, the sample was loaded onto the Ni-NTA column for contaminant and tag removal. The flow through containing the purified protein was collected, concentrated, and ran through an S75 size exclusion column (GE healthcare) in the NMR buffer (20 mM KPO_4_ pH 7.4, 50 mM KCl, and 0.5 mM EDTA. For PPARγ full-length and LBD the corresponding protein peak was collected and stored at -80 °C prior to use. For PPARγ AF1 the corresponding protein peak was collected, concentrated to 5mL and boiled at 70°C for 10 min. The sample was centrifuged at 4000 rpm for 15 min, filtered with a 0.2µm filter and loaded into a Q column (GE healthcare) to remove suspected lingering proteases and DnaK. The column was eluted with a 1M potassium chloride gradient where the most prominent peak corresponds to our protein of interest. The peak was collected and dialysed overnight in the NMR buffer. The next day, the protein was concentrated and stored at -80°C. All purified proteins were verified by SDS-PAGE as >95% pure and 1mM TCEP was added to all the buffers during PPARγ full-length purification.

### CD spectroscopy

Circular dichroism (CD) data of PPARγ AF-1 and LBD were collected using a Jasco J-815 CD Spectropolarimeter using a buffer containing 10 mM potassium phosphate and 50 mM potassium fluoride. An average of 3 scans were recorded per measurement using a scan speed of 100 nm/min at room temperature (∼23 °C) between a spectral range of 190-260 nm using a 1-mm optical bandwidth. Thermal unfolding curves were obtained by increasing the temperature from 20-80 °C with measurements recorded at 222 nm and 200 nm.

### Sortase A-mediated protein ligation

Purified Sortase 4M enzyme ^38^ was used to ligate ^15^N-labeled AF-1 (residues 1-135) + LPITG + 6xHis-tag protein (60 µM final concentration) to unlabeled Gly-DBD-hinge-LBD protein (residues 136-505; 60 µM final concentration); and unlabeled AF-1-DBD-hinge + LPITG + 6xHis-tag protein (60 µM final concentration) to ^15^N-labeled Gly-LBD (residues 231-505) protein. The ligation was performed in a buffer containing 50 mM TrisHCl (pH 7.8), 5 mM CaCl_2_, 100 mM NaCl, 0.2 mM TCEP, and 5 µM Sortase4M (final concentration). After 30 min, the ligation product, full-length PPARγ containing an LPITG insert between the ligated sequences, was purified using a HisTrapFF column washed with a buffer containing 40 mM phosphate buffer (pH 7.4), 500 mM KCl, 15 mM imidazole, and 1 mM TCEP and eluted in a similar buffer containing 500 mM imidazole. To remove the 6xHis-tag, TEV protease was added to the ligated sample and incubated overnight at 4 °C in a buffer containing 40 mM potassium phosphate (pH 7.4), 200 mM KCl, 0.5 mM EDTA, and 1 mM TCEP. Protein was reloaded onto the HisTrapFF column and the unbound fraction was collected and concentrated with a 30 kDa Amicon concentrator to 11.6 mg/mL (200 µM).

### MTSSL nitroxide spin labeling for PRE NMR studies

MTSSL labeling of proteins was performed following a published protocol ^30^. Briefly, proteins were concentrated to ∼300-500 µM in 1 mL, reduced with 1.25 mM dithiothreitol (DTT), and passed through a Zeba desalting column (Thermo Scientific) that had been equilibrated with NMR buffer; the eluate was collected, wrapped in aluminum foil, and supplemented with NMR buffer containing 10 molar equivalents of MTSSL (Cayman Chemical #16463) from a stock solution in 100% DMSO-d_6_. The labeling reaction was set up at room temperature (∼23 °C) for 15 min with gentle mixing followed by an overnight (∼18 hrs) incubation after addition of another 10 molar equivalents of MTSSL. The following day, the MTSSL-labeled protein was concentrated to 400 µL and dialysed for 16 hrs at 4 °C in the dark to remove unconjugated MTSSL.

### NMR spectroscopy

NMR experiments were acquired at 298 K (unless indicated otherwise) on a Bruker 700 MHz NMR instrument equipped with a QCI-P cryoprobe or a Bruker 600 MHz NMR instrument equipped with a TCI-H/F cryoprobe. NMR samples were prepared in NMR buffer (50 mM potassium phosphate, 20 mM potassium chloride, 1 mM TCEP, pH 7.4, 10% D_2_O) and typically contained 100-600 µM of the isotopically labeled component (^13^C and/or ^15^N); no significant chemical shift differences were observed within this concentration range. 3D NMR experiments used for AF-1 chemical shift assignment were collected using a 300 µM ^13^C,^15^N-labeled AF-1 sample with an N-terminal 6xHis+TEV tag and included HNCO, HN(CO)CA, CBCANH, CBCA(CO)NH, and CC(CO) NH. 2D NMR experiments included 2D [^1^H,^15^N]-HSQC, 2D [^13^C,^15^N]-(HACA)CON ^75^, and ZZ-exchange measurements via 2D [^1^H,^15^N]-HSQC for the AF-1 at different temperatures; and [^1^H,^15^N]-TROSY-HSQC of the LBD with or without 2 molar equivalents of rosiglitazone (Cayman Chemical #71740). 1D [^1^H]-NMR spectra were collected using natural abundance proteins that were not isotopically labeled. 3D [^1^H,^15^N,^1^H]-NOESY-HSQC and [^1^H,^15^N,^1^H]-TOCSY-HSQC experiments were used to aid in the transfer AF-1 assignments to mutant forms using the minimal chemical shift perturbation method ^76^ and residue-specific chemical shift trends from the BioMagResBank (BMRB) ^77^. Titration experiments for 2D NMR chemical shift structural footprinting were performed using 100 or 200 µM of the isotopically labeled component and the indicated equivalent of unlabeled component. PRE NMR experiments were collected in the absence or presence of 5x molar excess of sodium ascorbate to reduce the MTSSL nitroxide spin label; I_PRE_ values were calculated from the ratio of peak intensities (I_para_/I_dia_) as previously described ^30^. NMR data were collected using Bruker Topspin (v3.2) software, and processed/analyzed using NMRFx and NMRViewJ ^78,79^.

### Chemical crosslinking of protein samples

Full-length PPARγ and PPARγ ΔAF-1 (residues 136-505) were diluted to 10 µM in a buffer containing 20 mM potassium phosphate (pH 8.0), 150 mM KCl, 0.5 mM sodium citrate, and 10% glycerol. DSSO crosslinker (ThermoFisher #A33545) was freshly dissolved in DMSO to a final concentration of 75 mM and added to the protein solution at a final concentration of 1.5 mM. The reaction was incubated at 25 °C for 1 hr and then quenched by adding Tris buffer (pH 8.0) to a final concentration of 50 mM and incubating an additional 10 min at 25 °C. Control reactions were performed in parallel without adding the DSSO crosslinker. All crosslinking reactions were carried out in three replicates. Crosslinked samples were confirmed by SDS-PAGE and Coomassie staining along with negative control samples that were not treated with DSSO. Samples were separately pooled, precipitated using acetone, and dried protein pellets were resuspended to 12.5 µL in a buffer containing 50 mM ammonium bicarbonate (pH 8.0) and 8 M urea. ProteaseMAX (Promega, V5111) was added to the resuspended samples to a final concentration of 0.03%, the solutions were mixed on an orbital shaker operating at 1000 rpm for 15 min, and then 87.5 µL of 50 mM ammonium bicarbonate (pH 8.0) was added. Samples were digested for 4.5 hrs using trypsin added at a ratio of 1:170 (w/w trypsin:protein) at 37 °C then subsequently digested for 18 hrs using chymotrypsin at a ratio of 1:85 (w/w chymotrypsin:protein) at 25 °C. The resulting peptides were acidified to 0.67% trifluoroacetic acid (TFA) and then desalted using C18ZipTip (Millipore cat no. ZTC18 5096). Dried peptides were frozen, stored at -20°C, and resuspended in 10 µL of 0.1% TFA in water prior to LC-MS analysis.

### Chemical crosslinking mass spectrometry (XL-MS)

500 ng of sample was injected (triplicate injections for both control and crosslinked samples) onto an UltiMate 3000 UHP liquid chromatography system (Dionex, ThermoFisher). Peptides were trapped using a µPAC C18 trapping column (PharmaFluidics) using a load pump operating at 20 µL/min. Peptides were separated on a 200 cm µPAC C18 column (PharmaFluidics) using the following gradient: 5% Solvent B for 70 min, 30% Solvent B from 70 to 90 min, 55% Solvent B from 90 to 112 min, 97% Solvent B from 112 to 122 min, and 5% Solvent B from 120 to 140 min, at a flow rate of 800 nL/min. Gradient Solvent A contained 0.1% formic acid, and Solvent B contained 80% acetonitrile and 0.1% formic acid. Liquid chromatography eluate was interfaced to an Orbitrap Fusion Lumos Tribrid mass spectrometer (ThermoFisher) through a Nanospray Flex ion source (ThermoFisher). The source voltage was set to 2.5 kV, and the S-Lens RF level was set to 30%. Crosslinks were identified using a previously described MS2-MS3 method with slight modifications ^80^. Full scans were recorded from m/z 350 to 1,500 at a resolution of 60,000 in the Orbitrap mass analyzer. The AGC target value was set to 4×10^5^, and the maximum injection time was set to 50 ms in the Orbitrap. MS2 scans were recorded at a resolution of 30,000 in the Orbitrap mass analyzer. Only precursors with a charge state between 3 and 8 were selected for MS2 scans. The AGC target was set to 5×10^4^, a maximum injection time of 54 ms, and an isolation width of 1.6 m/z. The CID fragmentation energy was set to 25%. The two most abundant reporter doublets from the MS2 scans with a charge state of 2–6, a 31.9721 Da mass difference ^81^, and a mass tolerance of ±10 ppm were selected for MS3. The MS3 scans were recorded in the ion trap in rapid mode using HCD fragmentation with 35% collision energy. The AGC target was set to 20,000, and the maximum injection time was set for 150 ms and the isolation width to 2.0 m/z.

To identify crosslinked peptides, Thermo .Raw files were imported into Proteome Discoverer 2.5 (ThermoFisher) and analyzed via the XlinkX algorithm ^82^ using the MS2_MS3 workflow with the following parameters: MS1 mass tolerance, 10 ppm; MS2 mass tolerance, 20 ppm; MS3 mass tolerance, 0.6 Da; digestion, trypsin-chymotrypsin with ten missed cleavages allowed; minimum peptide length of five amino acids; and DSSO (K, S, T, Y). The XlinkX/PD Validator node was used for crosslinked peptide validation with a 5% false discovery rate (FDR). Identified crosslinks were further validated and quantified using Skyline (version 21.1) ^83^ using a previously described protocol ^84^. Crosslink spectral matches found in Proteome Discoverer were exported and converted to the sequence spectrum list format using Excel (Microsoft). Crosslink peak areas were assessed using the MS1 full-scan filtering protocol for peaks within 10 min of the crosslink spectral match identification. Peak areas were assigned to the specified crosslinked peptide identification if the mass error was within 10 ppm of the theoretical mass, if the isotope dot product was greater than 0.9, and if the peak was not found in the non-crosslinked negative control samples. The isotope dot product compares the distribution of the measured MS1 signals against the theoretical isotope abundance distribution calculated based on the peptide sequence. Its value ranges between 0 and 1, where 1 indicates a perfect match ^85^. Pairwise comparisons were made using the “MSstats” package ^86^ implemented in Skyline to calculate relative fold changes and significance. Significant change thresholds were defined as a log2 fold change ± 2 and -log10 p-value greater than 1.3 (i.e., a p-value less than 0.05). The visualization of proteins and crosslinks was generated using xiNET ^87^.

### Fluorescent labeling of protein samples for smFRET

Purified PPARγ AF-1 D33C+Q121C mutant protein at 50 µM was labeled with Alexa Fluor 488 and Alexa Fluor 647 maleimide dyes (Thermo Fisher) in 20 mM sodium phosphate pH 7.2, 50 mM NaCl, 1 mM TCEP using a 5 mM dye stock. Six substoichiometric additions of the dyes were made to the protein construct over 3 hrs for a final three-fold molar excess of each dye vs protein. The labeled sample was passed twice through Zeba desalting columns (Thermo Scientific) equilibrated in the same buffer to remove excess fluorophores.

### Single-molecule FRET (smFRET)

smFRET was performed using a two-color approach with a confocal setup where fluorescently labeled proteins diffuse freely. To record smFRET data, the labeled protein at 15 µM was diluted 100,000-fold into the buffer (10 mM sodium phosphate with specified pH values and NaCl concentrations) reaching a final concentration of approximately 150 pM. 300 µL of the diluted sample were deposited into Tween-20-coated (10% Tween 20, Sigma) coverslips (Nunc Lab-Tek Chambered Coverglass, Thermo Fisher Scientific). Background samples were prepared similarly but without protein. Fluorescence bursts were recorded over 3 hrs at 22 °C on a homebuilt multiparameter fluorescence detection microscope with pulsed interleaved excitation (MFD-PIE) as established elsewhere with minor modifications ^32^. Emission from a pulsed 483-nm laser diode (LDH-D-C-485, PicoQuant) was cleaned up (Semrock, FF01-482/25-25), emission from a 635-nm laser diode (LDH-D-C-640, PicoQuant) was cleaned up (Semrock, FF01-635/18-25), and both lasers were alternated at 30 MHz using a waveform generator (Keysight), a picosecond delayer (Micro Photon Devices) connected to the laser drivers (PDL 800-D). The red laser was delayed by ∼20 ns with respect to the blue laser. Linear polarization was cleaned up (Glan-Taylor Polarizer, Thorlabs, GT10-A) and the red and blue light were combined into a single-mode optical fiber (kineFlex, Point Source) before the light (100 µW of 483 nm light and 75 µW of 635 nm light) was reflected into the back port of the microscope (Axiuovert 200, Zeiss) and to the objective (C-APOCHROMAT, 40x/1.2 W, Zeiss). Sample emission was transmitted through a polychroic mirror (Chroma, ZT488/640rpc), focused through a 75-mm pinhole and spectrally split (Semrock FF593-Di03-25×36). The blue range was filtered (Semrock, FF03-525/50-25), and polarization was split (PBS101, Thorlabs) into parallel and perpendicular channels. The red range was also filtered (Semrock, FF01-698/70-25), and polarization was split (PBS101, Thorlabs). Photons were detected on four avalanche photodiodes (SPCM AQR 14, PerkinElmer Optoelectronics, for the green parallel and perpendicular channels and for the red parallel channel and SPCM AQRH 14, Excelitas, for the red perpendicular channel), which were connected to a time-correlated single-photon counting (TCSPC) device (MultiHarp 150N, PicoQuant). Signals were stored in 12-bit first-in-first-out (FIFO) files. Microscope alignment was carried out using fluorescence correlation spectroscopy (FCS) on freely diffusing ATTO 488-CA and ATTO 655-CA (ATTO-TEC). Instrument response functions (IRFs) were recorded one detector at-a-time in a solution of ATTO 488-CA or ATTO 655-CA in near-saturated centrifuged potassium iodide at a 25-kHz average count rate for a total of 25 × 106 photons. Macro Time-dependent microtime shifting was corrected for two (blue/parallel and red/perpendicular) of four avalanche photodiodes (APDs) using the IRF data as input. Data were analyzed using PIE Analysis with Matlab (PAM) software ^88^ using standard procedures for MFD-PIE smFRET burst analysis ^89,90^. Signals from each TCSPC routing channel (corresponding to the individual detectors) were divided in time gates to discern 483-nm excited FRET photons from 635-nm excited acceptor photons. A two-color MFD all-photon burst search algorithm using a 500-µs sliding time window (minimum of 100 photons per burst, minimum of 5 photons per time window) was used to identify single donor- and/or acceptor–labeled molecules in the fluorescence traces. Double-labeled single molecules were selected from the raw burst data using a kernel density estimator (ALEX-2CDE ≤ 15) that also excluded other artifacts ^91^. Sparse slow-diffusing aggregates were removed from the data by excluding bursts exhibiting a burst duration > 12 ms. By generating histograms of E versus measurement time, we corroborated that the distribution of E was invariant over the duration of the measurement. Data was corrected in this order to obtain the absolute stoichiometry parameter S and absolute FRET efficiency E: background subtraction, donor emission crosstalk correction, acceptor direct excitation correction and relative detection efficiency correction. To obtain the relative detection efficiency correction factor (γ), the center values of the E-S data cloud for each protein were estimated manually, plotted in an E vs. 1/S graph, and the data were fit to the following equation where Ω is the intercept and Σ the slope of the linear fit: γ = (Ω -1)/(Ω + Σ -1)

### Luciferase reporter assays

HEK293T cells (ATCC #CRL-11268) were seeded in a white 96-well plate at 15,000 cells/mL per well. The following day, cells were transfected using Lipofectamine 2000 (Thermo Fisher Scientific) and Opti-MEM with full-length PPARγ2 wild-type or P37A/F38A/W39A/P40A (PFWP-to-AAAA), or empty vector control (pcDNA3.1) expression plasmids (50 ng/well) along with pGL3-synthetic-PPRE-Luc2 (150 ng/well) incubated for 18 hrs at 37 °C, 5% CO_2_. The media was aspirated without disturbing the cells then replaced with media supplemented with either 1 µM Rosiglitazone or the same volume of 100% DMSO and incubated 18 hrs at 37 °C, 5% CO_2_. The cells were harvested for luciferase activity quantified using Britelite Plus (Perkin Elmer; 25 µL) on a Synergy Neo plate reader (Biotek). Data were plotted as mean ± s.d. in GraphPad Prism; statistics performed using two-way ANOVA with Tukey multiple comparisons analysis and are representative of ≥2 independent experiments.

### Computational analyses

Disordered structural predictions were performed with ODiNPred ^92^ using the human PPARγ2 isoform (505 residues). AF-1 secondary structure propensity calculation was performed with SSP ^28^ using AF-1 Ca and Cp NMR chemical shift assignments. The net charge per residue of the AF-1 sequence was calculated using localcider (https://pappulab.github.io/localCIDER/) ^93^. Predicted PRE profiles from an extended AF-1 conformational ensemble (n=10,000 structures) with no long-range contacts were calculated using flexible-meccano ^31^. Colabfold ^94^ was used to generate AlphaFold 2 ^18^ models, which were similar to the model obtained from the AlphaFold Protein Structure Database (https://alphafold.ebi.ac.uk/) that was used herein ^95^; AlphaFold 3 models were generated using the AlphaFold Server (https://alphafoldserver.com/) ^44^

## ACKNOWLEDGMENTS

This work was supported by National Institutes of Health (NIH) grants R01DK124870 (to D.J.K.) from the National Institute of Diabetes and Digestive and Kidney Diseases (NIDDK/DK); R01AG071332 (to P.R.G.) from the National Institute of Aging (NIA/AG); and R35GM130375 (to A.D.) from the National Institute of General Medical Sciences (NIGMS/ GM). The purchase of the 600 MHz NMR instrument at Scripps Florida/ UF Scripps was supported in part by NIH grant S10OD021550 from the NIH. Office of the Director. S.M. and C.C.W. were supported by NIH/ NIDDK (F31DK127643 and F31DK134167). K.-T.K. was supported by a Farris Foundation Fellowship. D.S. was supported by a postdoctoral fellowship from the Belgian American Educational Foundation. This content is solely the responsibility of the authors and does not necessarily represent the official views of the NIH or NIDDK.

## DATA AVAILABILITY

NMR chemical shift assignments of human PPARγ2 AF-1 (residues 1-135) have been deposited in the Biological Magnetic Resonance Data Bank (BMRB 51507). XL-MS data of full-length PPARγ and ΔAF-1 PPARγ have been deposited in the Proteomics Identification Database (PRIDE PXD035304). datasets generated and/or analyzed during the study are available from the corresponding author on reasonable request.

## AUTHOR CONTRIBUTIONS

S.M. and D.K. conceived and designed the research. S.M., P.M.T., B.M, X.Y., C.C.W. and R.B. prepared samples, performed experiments, and analyzed data. J.B. created mutant plasmids. J.L. and E.K. provided reagents and expertise. D.S. and A.A.D. performed and analyzed smFRET. K.-T.K., T.S.S., and P.R.G. performed and analyzed XL-MS. D.K. supervised the research and analyzed data along with the coauthors. S.M. and D.K. wrote the manuscript with input from all authors who approved the final version. The authors declare no conflicts of interest in the completion of this study.

