## Supplementary Figures S1-S2 and Table S1 for "Structural basis of interdomain communication in PPARγ"

[1] Skaggs Graduate School of Chemical and Biological Sciences at Scripps Research, Jupiter, United States
[2] Department of Integrative Structural and Computational Biology, Scripps Research and The Herbert Wertheim UF Scripps Institute for Biomedical Innovation & Technology, Jupiter, Florida, United States
[3] Department of Biochemistry, Vanderbilt University, Nashville, Tennessee, USA
[4] Department of Molecular Medicine, Scripps Research and The Herbert Wertheim UF Scripps Institute for Biomedical Innovation & Technology, Jupiter, Florida 33458, USA
[5] Department of Integrative Structural and Computational Biology, Scripps Research, La Jolla, California 92037, USA
[6] Structural Biophysics Laboratory, Center for Structural Biology, Center for Cancer Research, National Cancer Institute, Frederick, Maryland 21702, USA
[7] Center for Molecular Medicine, University Medical Center Utrecht, Utrecht University, Utrecht, The Netherlands
[8] Center for Structural Biology, Vanderbilt University, Nashville, Tennessee, United States
[9] Vanderbilt Institute of Chemical Biology, Vanderbilt University, Nashville, Tennessee, United States
[10] Center for Applied AI in Protein Dynamics, Vanderbilt University, Nashville, Tennessee, United States
[#] Co-first authors

File contains:

- Supplementary Figures S1–S2
- Supplementary Table S1

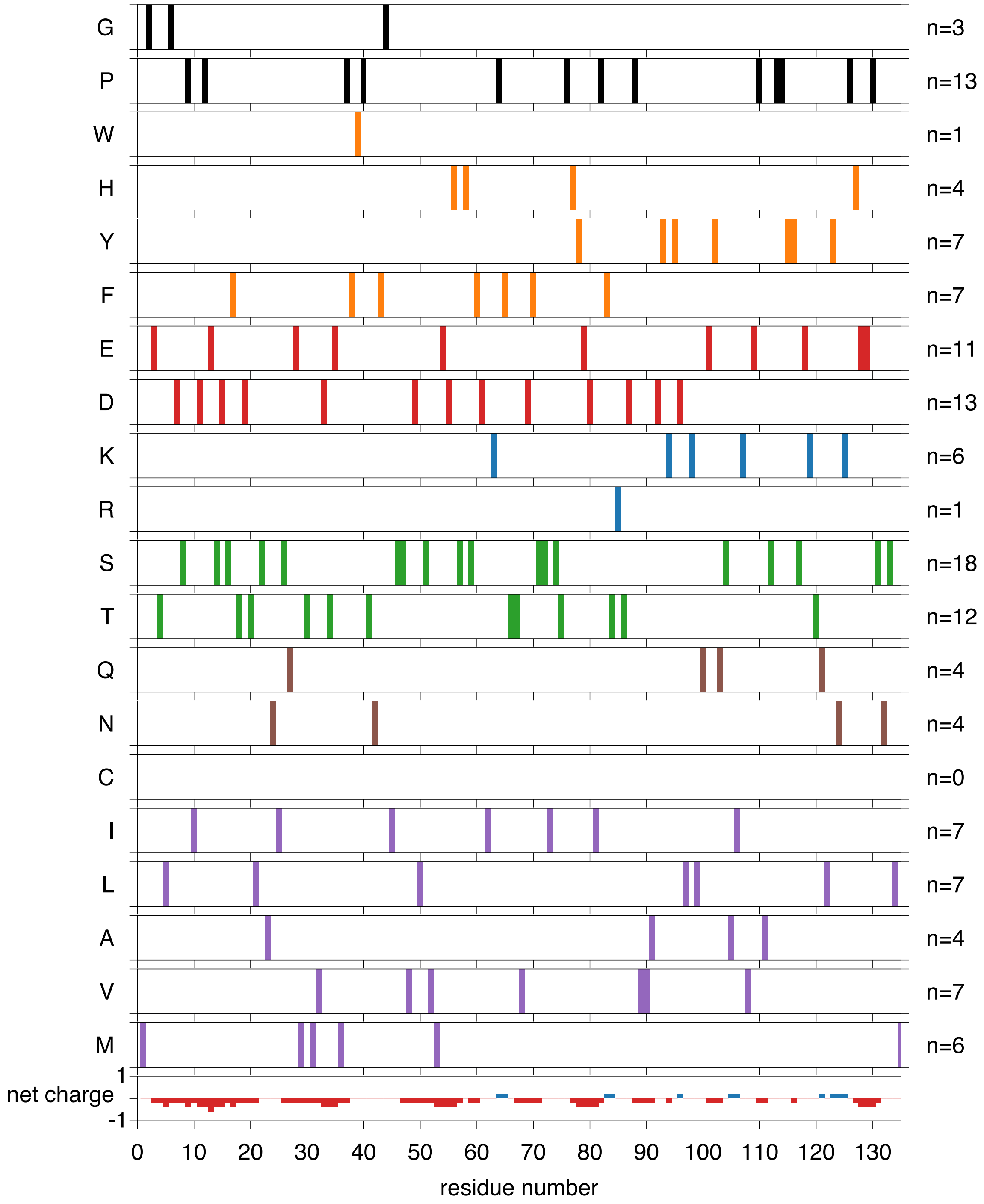


**Supplementary Figure S1.** Amino acid composition and net charge properties of the PPARγ AF-1. Residues are colored by property (purple, short/medium chain hydrophobic; brown, amide-containing; green, polar; blue, positively charged; red, negatively charged; orange, aromatic; black, other. Number of residues present in the AF-1 sequence is noted on the right. The net charge calculator ranges from 1 (positively charged) to -1 (negatively charged).

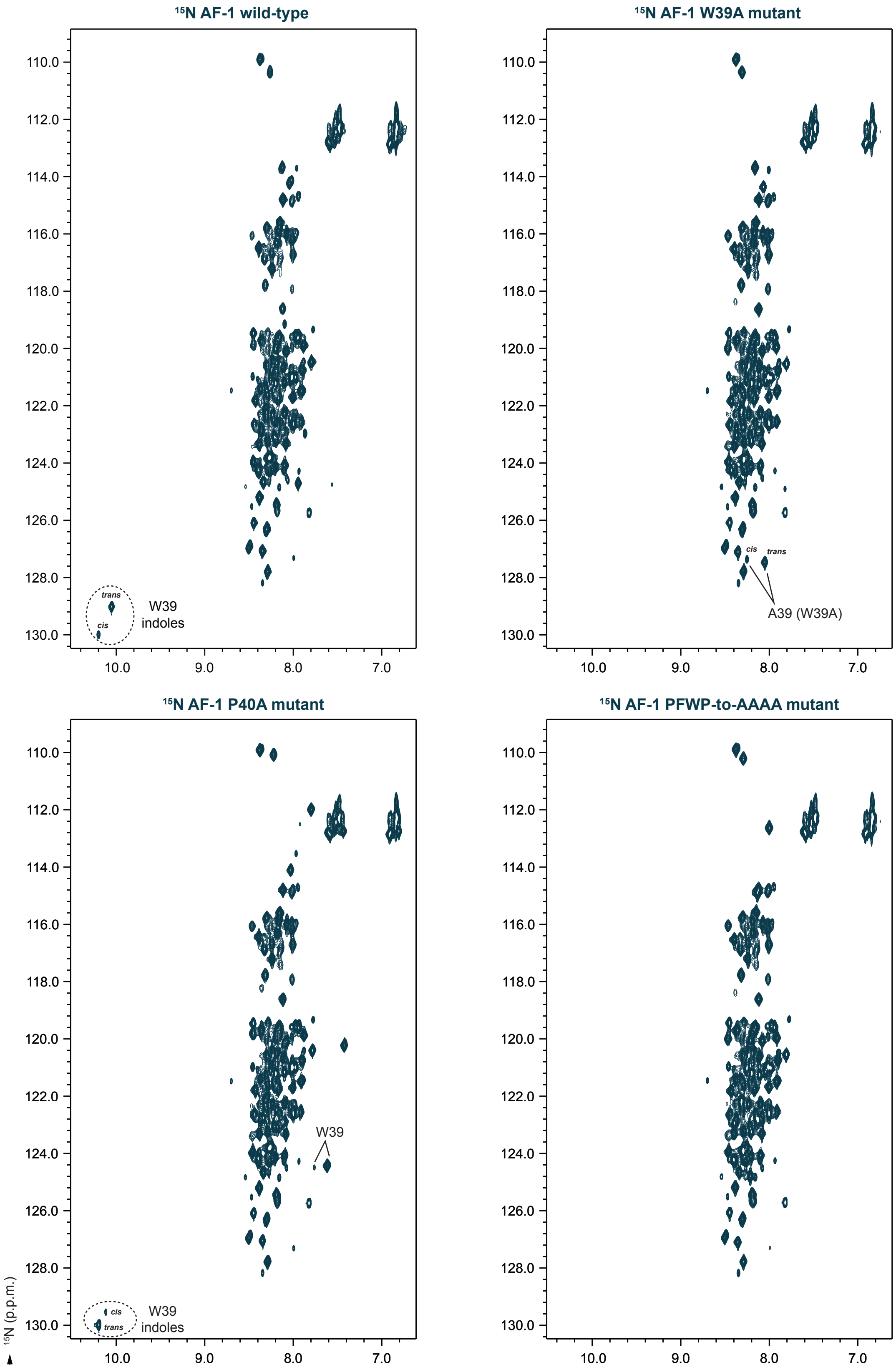


**Supplementary Figure S2.** 2D [$^{1}$H,$^{15}$N]-HSQC NMR spectra of (**A**) $^{15}$N-labeled PPARγ2 AF-1 and overlays of wild-type to mutant AF-1 constructs including (**B**) W39A, (**C**) P40A, and (**D**) PFWP-to-AAAA.

| sample, peaks | *trans* | *cis* |
|---|---|---|
| WT, W39 indoles | 0.84 | 0.16 |
| WT, W39 indoles | 0.89 | 0.11 |
| WT, W39 indoles | 0.83 | 0.17 |
| P40A, W39 indoles | 0.94 | 0.06 |
| P40A, W39 amides | 0.94 | 0.06 |
| W39A, A39 amides | 0.89 | 0.11 |

**Supplementary Table S1. Populations of the AF-1 cis and trans conformations from 2D [$^1$H,$^{15}$N]-HSQC NMR peak intensities.** Data for the WT W39 indole peaks (n=3) are representative of peak intensities measured from different samples during the course of our studies.
